# Comprehensive Analysis of Multi-Omics Vaccine Response Data Using MOFA and Stabl Algorithms

**DOI:** 10.1101/2025.09.30.679306

**Authors:** Aanya Gupta, Koji Abe, Holden T. Maecker

## Abstract

FluPRINT is a multi-omics dataset that measures donors’ protein expression and cell counts across various assays. Donors were also assigned a binary value (0 or 1), being labeled as high responders (1) if they had a fold change ≥ 4 of the antibody titer for hemagglutinin inhibition (HAI) from day 0 to day 28, and low responders otherwise (0). In this project, we used the MOFA and Stabl algorithms to analyze FluPRINT, estimate the population structure from the data, and identify the most important features for predicting response to the vaccine. The preprocessing of the dataset included removing repeat features, scaling by assay, and removing outliers. Since Stabl does not directly address missing values, features with high amounts of missing values were removed and the remaining were ignored. MOFA identified the top feature in structure extraction as IL neg 2 CD4 pos CD45RA neg pSTAT5. MOFA explains well the variance of the data while also choosing features that have good significance, as illustrated by their significant p-values (p < 0.05). Stabl found the top feature for explaining the outcome to be CD33- CD3+ CD4+ CD25hiCD127low CD161+ CD45RA+ Tregs, which matched the top result of previously published analysis. MOFA’s features achieved an AUROC of 0.616 (95% CI of 0.426-0.806), and Stabl’s achieved an AUROC of 0.634 (95% CI of 0.432-0.823). Our research addresses a key knowledge gap: understanding how these fundamentally different analytical approaches perform when analyzing the same complex dataset. Our exploration evaluates their respective strengths, limitations, and biological insights and provides guidance on using MOFA and Stabl to find the best predictive cell subsets and features for understanding large immunological multi-omics data. The code for this project can be found at https://github.com/aanya21gupta/fluprint.

## 1. Introduction

Technological advances now allow for multi-omics data, down to the single-cell level, and pave the way for genomic, transcriptomic, proteomic, and metabolomic profiling. With the rise of such highly complex, large amounts of immunological data, the potential to build a much more comprehensive and integrative biological analysis has grown (1). These analyses can be key tools for areas such as systems vaccinology, where large datasets on the immune states of individuals before and after vaccination are generated.

Before the rise of such datasets, it was hard to pinpoint specific cell subsets and cellular features that would be most important in predicting vaccine response. Now, several computational, machine learning-based approaches have aimed to use these datasets to either identify factors that drive differences in individual vaccine responses or create candidate biomarkers at much faster rate, accelerating the overall process of scientific discovery (Table 1). For example, the SIMON (Sequential Iterative Modeling OverNight) algorithm, an automated machine learning system, has been used to identify predictive biomarkers by comparing results from multiple algorithms and effectively handling missing data through subset creation (2). SIMON has successfully identified several predictive cell subsets and biomarkers associated with influenza vaccine response, demonstrating the potential of computational methods in biomarker discovery.

**Table 1.**
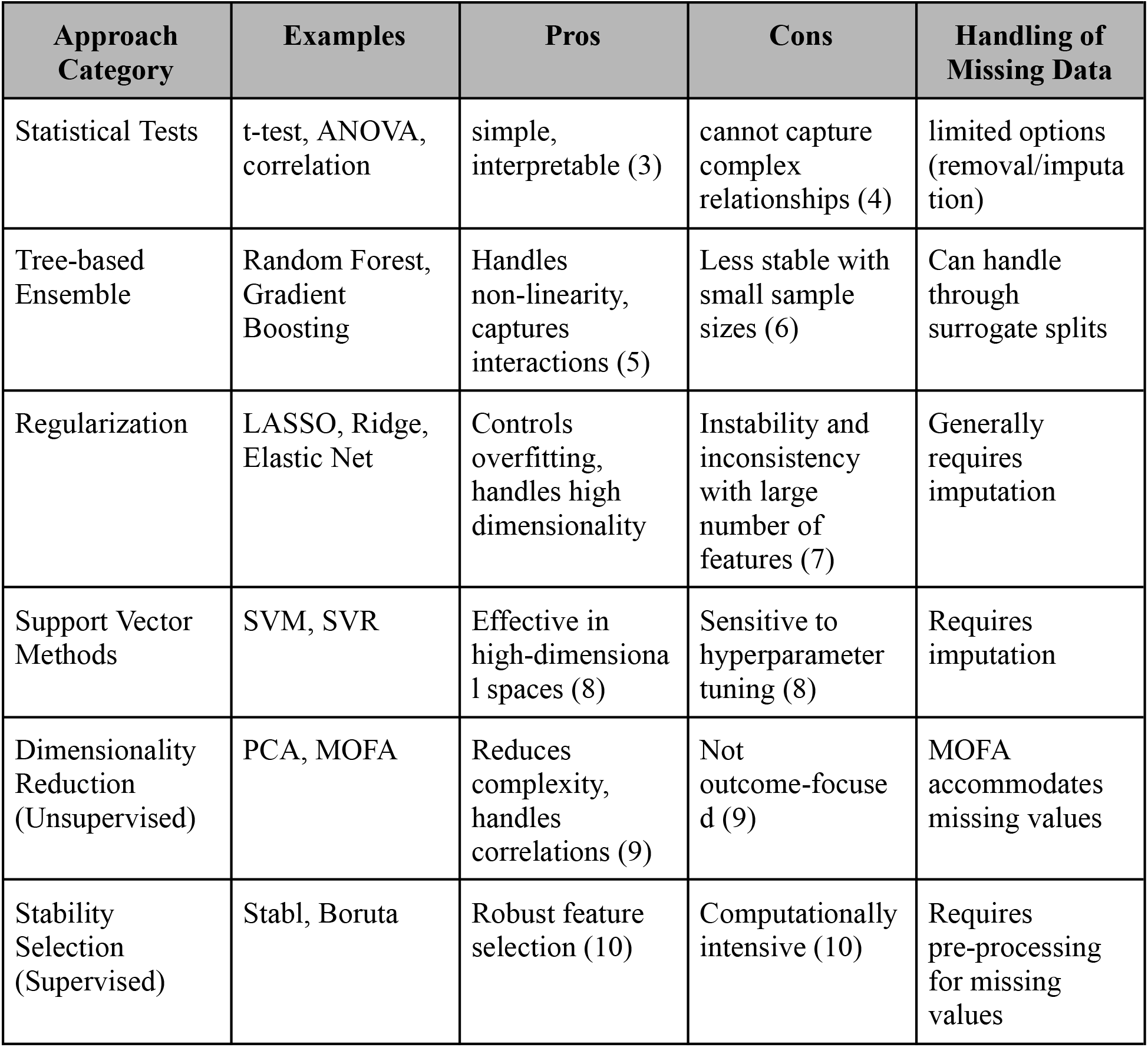
Common Methods of Feature Selection & Predictive Models.

However, there are many issues that come with the massive amounts of data available today, as seen in Table 1. High dimensionality relative to the number of samples can lead to overfitting. Ways of handling missing data and difference of units between assays are also variables that can yield large differences in findings. Missing values can derive from aggregation of studies with variations in the assays applied, a common situation in large longitudinal studies. Existing models can also suffer from lack of explainability and sparsity, which then also reduces interpretability in real-world context. These models face a ‘black box’ problem, where their complex internal workings make it hard to decipher their understanding of the complex relationships in both the proteomics and cell count data. They can also be extremely computationally intensive and highly sensitive to parameter choices. Thus, existing methods, including SIMON, often face challenges related to handling missing data and integrating heterogeneous multi-omics data, limiting their practical utility and biological insight.

In this study, we aim to address these limitations by exploring two complementary computational methods, Multi-Omics Factor Analysis (MOFA) and Stabl, to analyze the FluPRINT dataset. Our primary goal is to identify key biological features and cell subsets predictive of differential influenza vaccine responses (high vs. low responders). MOFA, or multi-omics factor analysis, discovers the principal sources of variation in multi-omics data sets by an unsupervised creation of a set of latent factors that capture biological and technical sources of variability, working well with multi-omics data (11). MOFA utilizes likelihoods in estimation, and can effectively deal with missing values by not including in the likelihood (with the assumption of data being missing at random). Stabl takes a different approach, as a supervised machine-learning based algorithm that identifies a very sparse, reliable set of predictive biomarkers using a unique threshold determined by the data and noise injection done by the algorithm (12).

MOFA and Stabl were designed especially to deal with these important and common problems that arise with other machine learning and statistical approaches. MOFA’s PCA-like unsupervised approach is not very sensitive to model parameters and provides for easy interpretability of results through simple correlations or weights (11). Stabl also addresses these issues, ensuring a robust process of feature selection using bootstrapping and ridge regularization (12). Logistic regression coupled with ridge regularization is uniquely equipped to handle feature selection due to its increased emphasis on the sparsity of the model. Due to its ElasticNet penalty, it also has the advantage of balancing between L1 and L2 regularization, encouraging a smaller number of features without making the model too dependent on a few features. Additionally, a direct methodological comparison between unsupervised and supervised approaches remains unexplored. Such a comparison is crucial for understanding how algorithmic selection impacts feature identification and interpretation in vaccine response studies. Here, we address this gap by implementing both MOFA (unsupervised) and Stabl (supervised) on the same dataset, evaluating their performance, concordance with previous findings, and unique contributions to understanding vaccine response predictors. Thus, in this paper, we explore these two complementary algorithms by analyzing the unique benefits and results of each on FluPRINT, an influenza vaccine dataset, to gain deeper biological insights and identify robust predictive biomarkers of influenza vaccine response.

## 2. Materials and Methods

### 2.1 Subjects, sample, and data collection

The FluPRINT dataset was created by combining the results of eight clinical studies from 2007 to 2015 (13). There were 740 individuals undergoing influenza vaccination (either IIV or LAIV) who had blood and serum samples taken at both baseline and post-vaccination timepoints (13). The original population had donors from ages 1-90 years, with a median age of 27 years, a distribution of 446 females and 294 males, and had a majority of Caucasians (13). Response was recorded as a binary value, with an individual considered a high responder (1) if they had a fold change greater than or equal to 4 in the antibody titer (HAI) from day 0 to day 28 and a low responder otherwise (0). Ideally, individuals in training data would be labeled as protected and unprotected following vaccination, but that cannot be the case for flu infection. Instead, hemagglutination inhibition (HAI) titer is very commonly used as a surrogate marker of protection and a fold-change (day 28/day 0 HAI) is often used to assess vaccine efficacy (eg., high and low responders based on 4-fold change relative to pre-vaccination titer) (14). Other factors that were recorded with each individual included gender, race, visit age, BMI, vaccine history, influenza history, cytomegalovirus (CMV) status, Epstein-Barr Virus (EBV) status, and statin use. The various assays included multiple cytokine assays (Luminex), hemagglutination inhibition assay, serological assays for CMV and EBV antibodies, phosphoepitope flow cytometry, and mass cytometry immunophenotyping, creating a heterogenous dataset with data from different assays, with both proteomics and cell count assays that also differed in units and thus ranges of their measurements.

For this project, the subset of individuals ranging from age 8-40 who received inactivated influenza vaccine (IIV) was taken, resulting in 187 donors (2). All assays were taken for each donor’s first visit, and this subset of the data was taken to minimize missing values (2). For this dataset, the “name” and “subset” columns were combined to create the features for MOFA and Stabl. Three assays were dropped, as documentation suggests that MOFA struggles to learning meaningful factors from assays with less than 15 features (11).

### 2.2 Pre-processing of data prior to model input

For both MOFA and Stabl analyses, the input data consisted of a data matrix with dimensions 162 × 3091, where each row represented one donor and each column represented a biological feature measured by one of the assays. The dataset included 162 donors selected from the original FluPRINT dataset. Each donor had multi-omics measurements taken at baseline (pre-vaccination) and post-vaccination timepoints. For the analyses presented here, we specifically used data from baseline measurements only. Demographic and clinical variables such as BMI, sex, age, vaccine history, influenza history, CMV and EBV status, and statin use were not included as input features. Supplementary Table 1 details the original assays and number of features for each.

MOFA takes as input the multi-omics data described above and outputs a set of latent factors, each representing a principal source of variation. The output from MOFA included factor loadings for each feature, indicating the contribution of each feature to each latent factor. To identify features potentially predictive of vaccine response, we correlated the latent factors identified by MOFA with the vaccine response outcome (high vs. low responders), and then selected the features with the highest loadings on the most strongly correlated factors.

The input to Stabl was the same multi-omics data matrix described above, along with the binary vaccine response outcome (high vs. low responders) for each donor, defined based on fold-change in antibody titers from pre- to post-vaccination. Stabl identified a sparse set of features most predictive of vaccine response by employing logistic regression with ridge regularization and bootstrapping to ensure robustness and stability of selected features. The output from Stabl included a sparse set of predictive features and predictive performance metrics (e.g., AUROC) for classifying high vs. low vaccine responders.

The various transformations that were applied to the data which was input into each algorithm are described below. Each assay used a single unit of measurement, allowing the data from each assay to be normalized consistently. Each assay used only one unit, allowing for the data to be scaled to a normal model, by assay. Outliers were then removed from each assay using the interquartile range method, with outliers being counted as data beyond 1.5 * IQR (interquartile range, data between the 25^th^ and 75^th^ percentile) of Q1 and Q3 (the 25^th^ and 75^th^ percentile of the data). This was done to prevent influential points that might have high leverage or be erroneous values from having a very large effect on the model. The Luminex assays (Human_Luminex 50, 51, and 62_63) were combined into one assay that only contained features that had been present in all three assays, while the rest were dropped. Afterwards, out of 16 assays, three were dropped for having fewer than 15 features. The only preprocessing that was different between the models was that Stabl required additional preprocessing. Features that had a fraction of missing values above a given threshold (40%) were dropped (Low Info Filter) and zero variance features were also removed, before imputing the remaining missing values using a KNN Imputer with three neighbors. The resulting eight assays, and number of features for each, are shown in Table 2.

**Table 2.** List of final assays (and corresponding number of features) for input data into MOFA and Stabl algorithms.

| Assay | Number of Features |
| --- | --- |
| CBC with Differential | 18 |
| Other_Luminex | 38 |
| Luminex_combined | 47 |
| Lyoplate_1 | 65 |
| CyTOF_phenotyping | 76 |
| Phospho_flow_cytokine_stim_(PBMC) | 274 |
| pCyTOF_(whole_blood)_pheno | 312 |
| pCyTOF_(whole_blood)_phospho | 2286 |

The assay Other_Luminex represents was performed only for one of the studies included in the FluPRINT dataset (study SLVP015 in 2007 using the Human 42-Plex Polystyrene Kit), and was preprocessed in the same way as the other Luminex assays.

### 2.3 Overview of MOFA, Multi-Omics Factor Analysis

MOFA, or Multi-Omics Factor Analysis, infers a PCA-like representation of the data through a few latent factors (11). It uses an unsupervised model to perform principal component analysis and learn latent factors from the data. To address the heterogeneity of the data due to multiple assays, MOFA allows for preprocessing and viewing of data and results by assay, allowing each assay to be scaled separately and its influence on each factor to be considered separately from other assays. The data is arranged in a matrix, where each row is a feature measured for a specific donor. If a feature is not measured for a sample, then that value in the matrix is considered missing. MOFA’s goal is then to use matrix algebra and machine learning to decompose the matrix into two matrices, which represent the relationships between the latent factors and the features and the factors and the samples.

### 2.4 Latent factor creation using MOFA

The model, given a set number of maximum factors, computes each factor as a linear combination of all the features. It does matrix factorization on the large matrix of all the data, where the structure of the data is specified in the prior distributions of the Bayesian model (11). ARD (automatic relevance determination) of the factors is done by sparsity priors, and the algorithm automatically drops any factors that explain less than 2% of the variance in the data (11). MOFA can handle missing values during model training, unlike some other methods which may require complete datasets. MOFA employs likelihoods in estimation, and missing data are not included in the likelihood calculations. These missing values are natively accounted for within the probabilistic framework of MOFA. Once the MOFA model is fit, missing values can be imputed by the MOFA pipeline. The resulting factors can then be further investigated using MOFA, to find the factors and corresponding features that are most related to the vaccine outcome.

### 2.5 Overview of Stabl

The Stabl algorithm, on the other hand, utilizes a different method for feature selection (12). Stabl inputs data as a matrix with each row representing one donor and each column representing a feature. Stabl takes random sampling of donors (with replacement) at a time, via bootstrapping and then on each subset of data, it uses an algorithm such as Lasso or ElasticNet to perform feature selection. To ensure the stability of these features, artificial noise (through artificial features) is added to the data, and then for varying values of lambda, the regularization parameter, the base model selects features out of this new, combined data. A feature is considered more robust, and more likely to be connected to outcome, if it is selected more times (in multiple samples). This process is repeated over values of lambda, a parameter which controls the amount of regularization of the base algorithm to ensure that the model is sparse while penalizing larger weights, which are the coefficients of the features in Stabl’s regression model. Features that have a probability of selection higher than a threshold are considered the most important features and then can be used for building a predictive model. The threshold is chosen as the threshold at the minimum of the FDP+ (false discovery proportion surrogate), which compares the number of artificial features injected to the number of selected actual molecular features, occurs. A Low Info Filter was also applied, which dropped features which had a percentage of missing values higher than a specified threshold before applying the feature selection process.

### 2.6 Criteria for significance of chosen features

For MOFA, all features had a weight for each factor which represented their correlation, or importance, to that factor. To extract the final subset of features for MOFA, the factor most correlated to outcome was found (based on correlation coefficient to outcome), and its most weighted features were taken as the only input for the final regression model. For Stabl, the features that surpassed the established threshold were considered significant and used exclusively in the model. Overall, the features selected by both MOFA and Stabl were then noted as significant to the outcome if a p-value < 0.05 was observed with a t-test for a significant difference in the means of the two independent samples of values.

### 2.7 Code and data availability

The code for this project is linked at https://github.com/aanya21gupta/fluprint. For this project, Python version 3.13.0 was used. Additionally, Mofapy2 version 0.7.2 and Stabl 1.0.1 were utilized.

## 3 Results

### 3.1 Application of MOFA model for all-relevant cellular predictors

The model was set to create 10 latent factors, training with 100 iterations in the ‘fast’ convergence mode. Ten factors were chosen as a starting value. The subsequent MOFA model had only 5 factors, indicating that more factors were not needed as they were dropped by the algorithm due to low variability explained. Fewer factors were not taken as this would make it harder to extract the most important features as the model would be forced to ignore potentially important variability to fit into a lesser number of factors. The final results were saved in a separate file to obtain the final model.

Figure 1 **(A)** displays the R^2^ values for each of the five factors separated by group. R^2^ is a statistical concept that measures level of correlation in the association between two variables; specifically, it is the percent of the variability in Ythat can be explained by the equation built to predict Yusing x. Factor 5 has the highest R^2^ value for group 0 and a negative value for group 1. This indicates that Factor 5 explains the most variance in group 0 but does not explain any of the variance in group 1, and thus is unlikely to explain outcome. Figure 1 **(B)** displays the average factor value for each group (outcome). Factor 1 was shown to have the sharpest difference in average factor value per group, having a positive average value for an outcome of 0 (0.085) and a negative average value for an outcome of 1 (-0.124). Figure 1 **(C)** demonstrates the correlation between the factors and the groups (outcome). Factor 1 had the strongest correlation (calculated Pearson correlation coefficient) to the outcome, with the highest positive correlation with an outcome of 0 (low responder) and highest negative correlation with an outcome of 1 (high responder).

**Figure 1.**
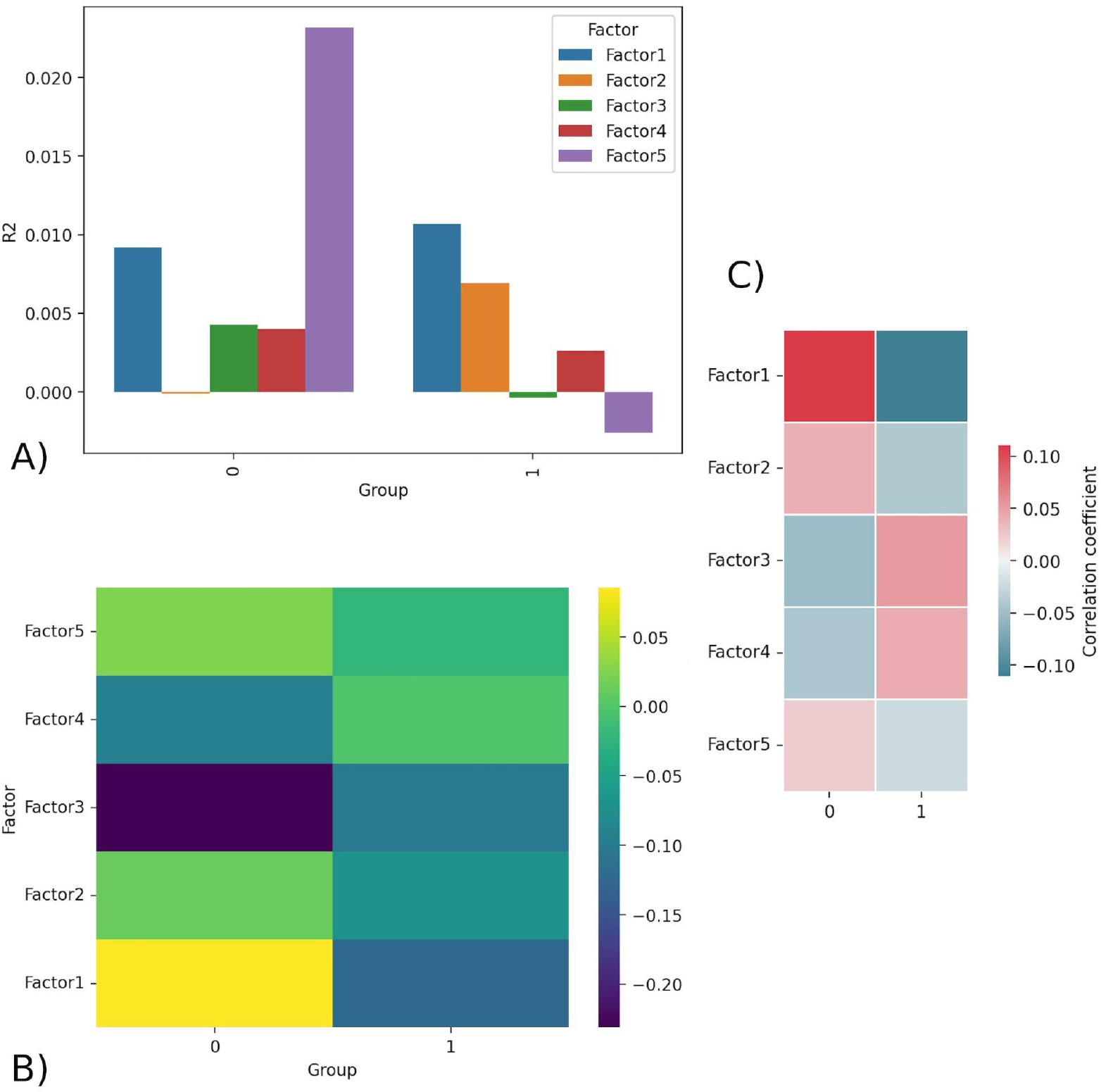
**A)** R^2^ barplot - R^2^ values for all factors for group 0 (low responder) and 1 (high responder). Factor 5 displays the highest R^2^ in group 0 and negative R^2^ in group 1, indicating it is likely unimportant in explaining outcome. **B)** Factors matrix - Average value of factor for group 0 and 1. Factor 1 has the largest difference in values, being strongly positive for group 0 (0.085) and strongly negative for group 1 (-0.124). **C)** Correlation - shows the Pearson correlation coefficient calculated between the continuous factor values (scores) for each donor and the binary outcome variable

Each factor is a linear, weighted combination of all features based on their importance in determining that factor. Therefore, the highest weighted features of Factor 1, the most correlated feature to outcome, were considered by MOFA to be the most highly correlated to response.

Table 3 displays the top ten features (and respective assays) of Factor 1 (ordered by magnitude of weight). Figure 2 displays the violin plots showing the distribution of these across outcomes along with if these two distributions are significantly different using a t-test for the means of two independent samples of values, indicating whether the feature actually differentiates outcome using p-value. The top weighted feature, IL neg 2 CD4 pos CD45RA neg pSTAT5 had a small p-value of 0.003 (assay: PBMC), while the most significant p-value (0.002) was that of IL neg 2 CD8 pos CD45RA neg pSTAT5 (assay: PBMC). The p-values were calculated using independent two-sample t-tests comparing the distributions of features and were unadjusted.

**Table 3.** Top 10 most weighted features of MOFA Factor 1, and therefore the most important features for determining outcome according to MOFA

| feature | subset | weight | assay |
| --- | --- | --- | --- |
| IL_neg_2_CD4_pos_CD45RA_neg_pSTAT5 | CD4+CD45RA-: pSTAT5 | 0.467 | PBMC |
| IL_neg_10_CD4_pos_CD45RA_neg_pSTAT1 | CD4+CD45RA-: pSTAT1 | 0.375 | PBMC |
| IL_neg_2_CD4_pos_pSTAT5 | CD4+: pSTAT5 | 0.356 | PBMC |
| IL_neg_10_CD4_pos_pSTAT1 | CD4+: pSTAT1 | 0.347 | PBMC |
| IL_neg_10_CD8_pos_CD45RA_pos_pSTAT1 | CD8+CD45RA-: pSTAT1 | 0.334 | PBMC |
| IFNa_B_cell_pSTAT1 | B cell: pSTAT1 | 0.316 | PBMC |
| IL_neg_10_CD8_pos_pSTAT1 | CD8+: pSTAT1 | 0.312 | PBMC |
| IL_neg_7_CD8_pos_CD45RA_neg_pSTAT5 | CD8+CD45RA-: pSTAT5 | 0.308 | PBMC |
| IL_neg_10_CD8_pos_CD45RA_neg_pSTAT1 | CD8+CD45RA-: pSTAT1 | 0.307 | PBMC |
| IL_neg_2_CD8_pos_CD45RA_neg_pSTAT5 | CD8+CD45RA-: pSTAT5 | 0.294 | PBMC |

**Figure 2.**
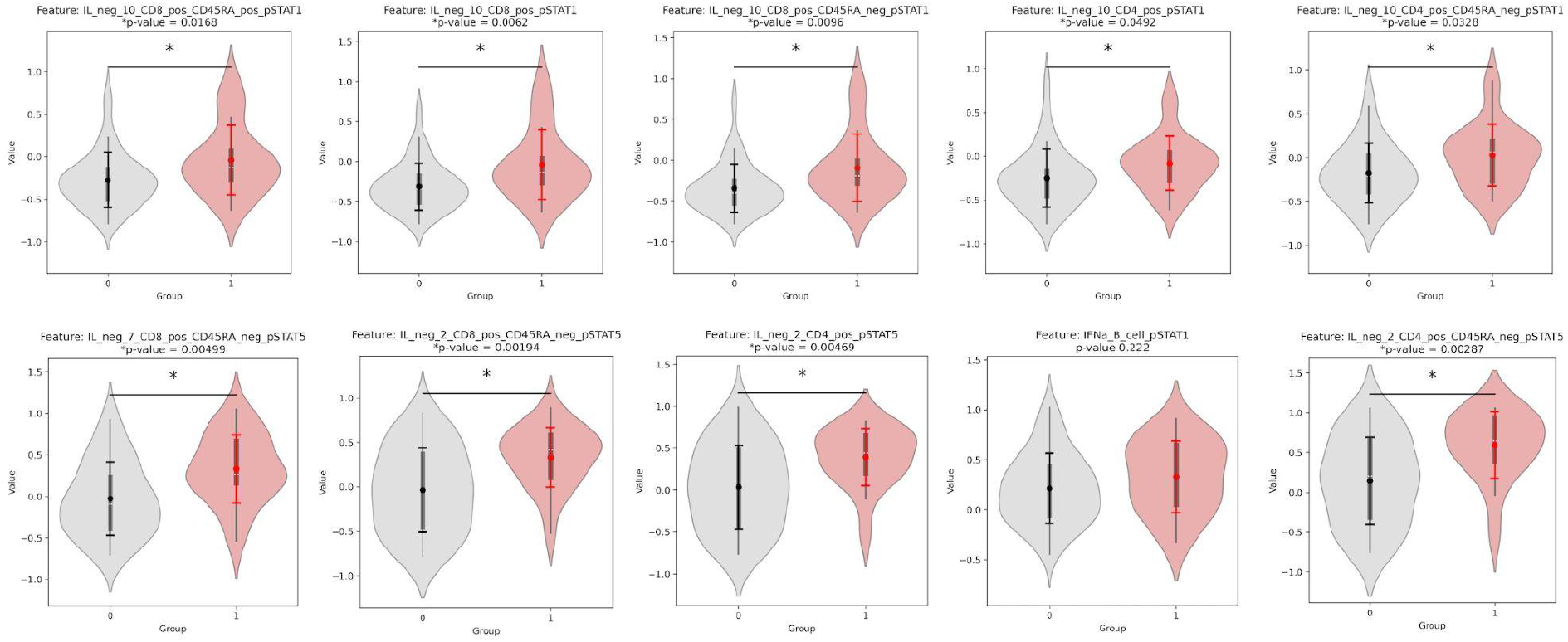
Violinplot distributions for group 0 vs. group 1 and p-values for a significant difference in distribution, for each of the top 10 weighted features of Factor 1. All features are from the PBMC assay, and their weights for Factor 1 are found in Table 3.

### 3.2 Predictive Modeling Using MOFA Top Features

To test the reliability of MOFA’s features in predicting response, we predicted outcome using only the top ten features that MOFA had found, since this number gave the best AUROC. These were the top ten most features with the highest magnitudes of weight in Factor 1. The data from only these factors was isolated from the dataset and then was split into a train-test split of 0.8:0.2. Any missing values were imputed using the KNN imputer with three neighbors, as it uses the similarity of nearby data points to account for the connectedness and intricate relationships between the complex data. The final predictive model was a logistic regression with an ElasticNet penalty, saga solver, and L1 to L2 regularization ratio of 0.5. The ROC curves for the training and testing data are shown below in Figure 3. MOFA’s top features achieved an AUROC of 0.616 on both the testing (95% CI of 0.426-0.806) and training data (95% CI of 0.521-0.711). The confidence interval of the AUROC in MOFA was calculated using a 95% confidence interval with the standard error of the AUROC calculated using the AUROC and the sample sizes of both classes (outcomes).

**Figure 3.**
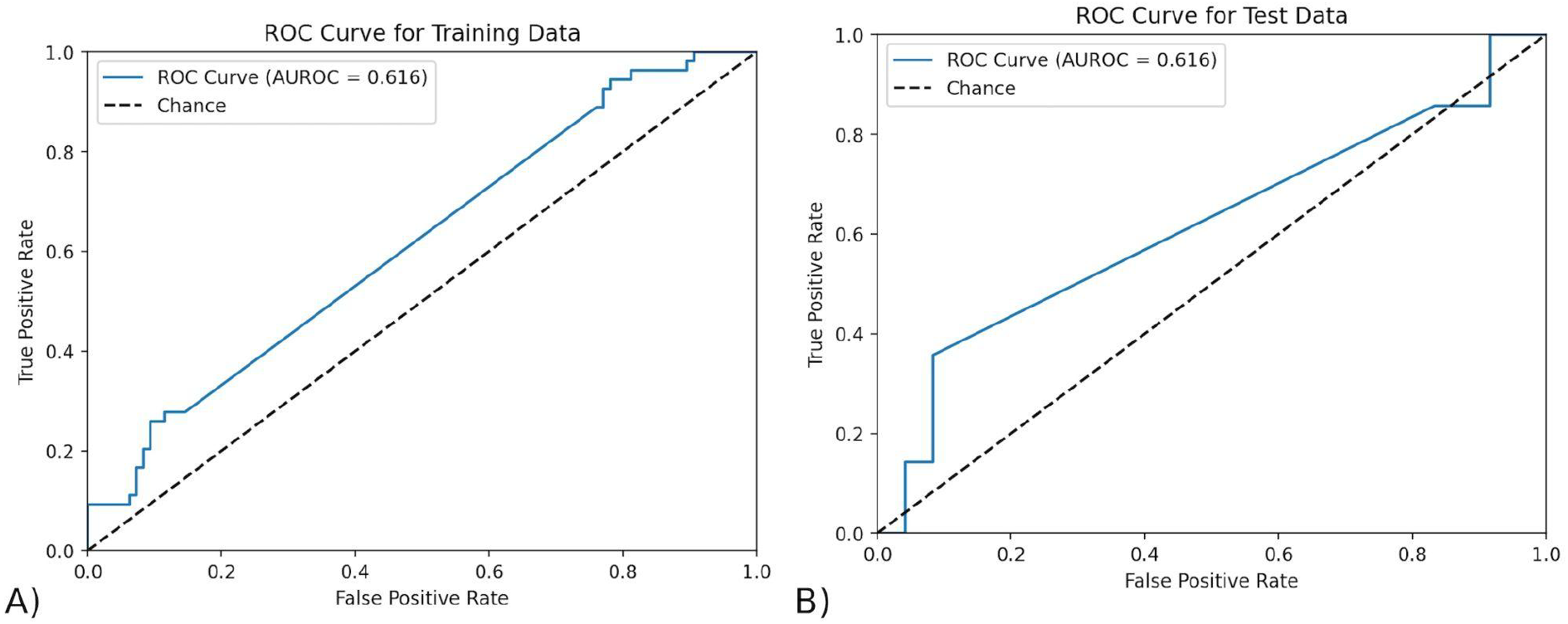
– The ROC curve of the model trained by MOFA features - **A)** The ROC curve on testing data. The AUROC of the testing data using exclusively the features found by MOFA is 0.616 (95% CI of 0.426-0.806) and training data. **B)** The ROC curve on training data. The AUROC of the training data using exclusively the features found by MOFA is 0.616 (95% CI of 0.521-0.711).

### 3.3 Application of Stabl model for all-relevant cellular predictors

For Stabl, after the zero variance features were removed and the Low Info Filter of 0.4 was applied, the remaining missing values were imputed with a KNN imputer that used three neighbors. The data was then scaled again, and split into train and test with 0.8:0.2 split. The Stabl algorithm was performed with a base estimator, or model, of a Logistic Regression with L1 penalty, liblinear solver, and balanced class weights. Balanced class weights ensured that the class frequencies (number of donors for each group) were taken into account when adjusting model weights. The Stabl class was run on the training data with 1000 bootstrap resamples and knockoff artificial features, which is noise injected into the input data as features that mimic those of the actual dataset. It tested out 20 lambdas, or level of overall regularization, automatically spaced out between 0 and 1. The same top features were found with a Logistic Regression with ElasticNet penalty, saga solver, and alpha (L1:L2 regularization ratio) of 0.5.

The top features for this version were CD14-CD33-CD3+ CD4+ CD25hiCD127low CD161+ CD45RA+ Tregs, CD14+ CD33+ monocytes, and CD4-CD33-CD3+ CD56+ NKT cells. These are decided as the features whose frequency of selection are greater than the FDP+ threshold (0.82), which is the threshold at which the rate at which false positives (Stabl-injected features) are chosen is minimized (Figure 4).

**Figure 4.**
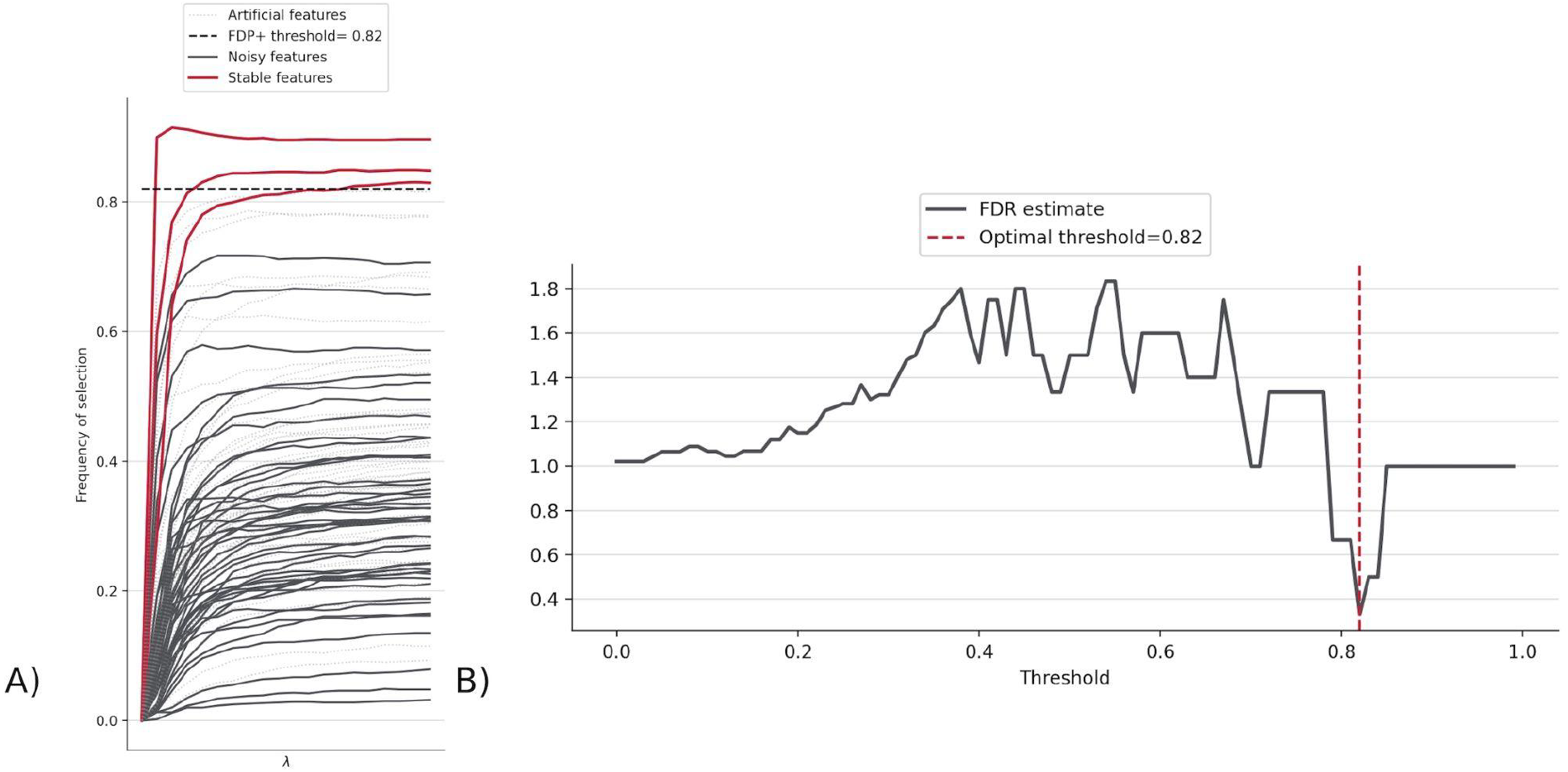
**A)** Stabl path showing the frequency of selection for each feature indicating how many features made it above FDP+ threshold (0.82). The three lines are features whose probability of being selected is higher than the threshold, and thus represent the top features found by Stabl. **B)** FDR graph, which displays the FDR estimate across different FDP+ thresholds, to determine the threshold where the chance of choosing a false positive/Stabl-injected feature (called the FDR estimate) is minimized, at 0.82 for FluPRINT.

CD14-CD33-CD3+ CD4+ CD25hi CD127low CD161+ CD45RA+ Tregs was given with all random states taken, with varying amounts of other features. Overall, the different top features found were: CD14-CD33-CD3+ CD4+ CD25hi CD127low CD161+ CD45RA+ Tregs, CD14+ CD33+ monocytes, CD4-CD33-CD3+ CD56+ NKT cells, and CD14-CD33-CD3+ CD4-CD8+ Non-naive CD8+ CXCR5+ TFH T cells. To identify the features that were significant predictors, we calculated the p-value to determine significant differences in values for donors with outcome of 0 versus donors with outcome of 1. The p-value for the top three features found with the initial random seed are shown below (Figure 5). CD14-CD3-CD3+ CD4+ CD25hi CD127low CD161+ CD45RA+ Tregs had an especially low p-value of approximately 0.0007.

**Figure 5.**
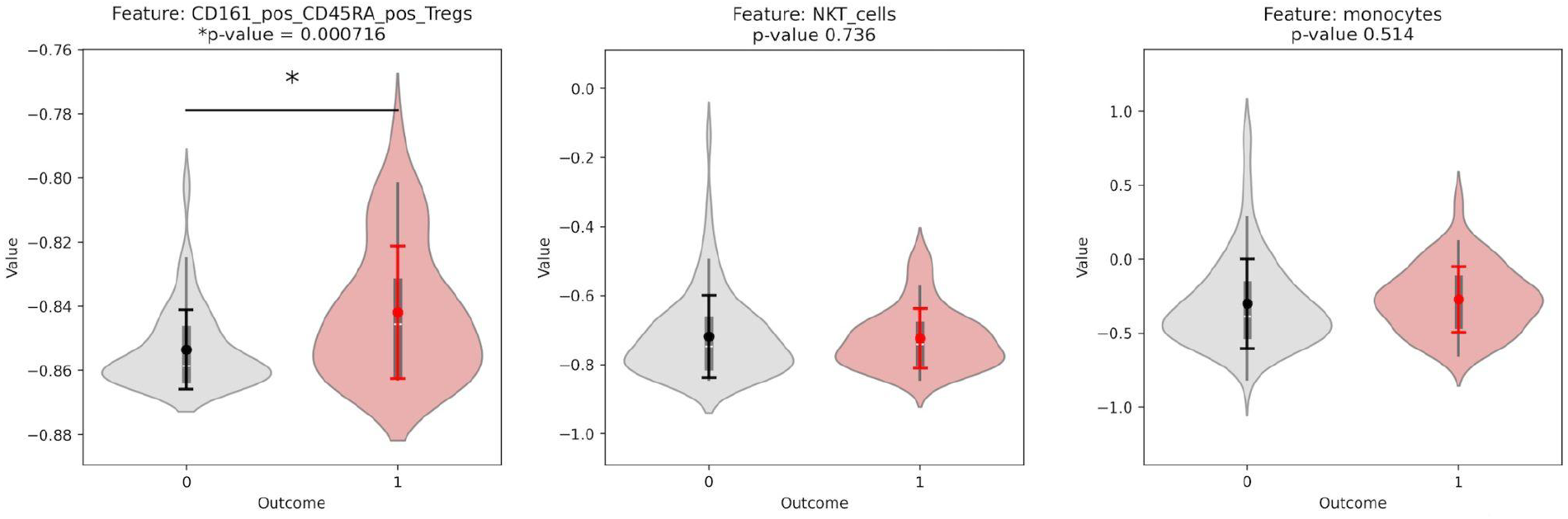
Violinplot distributions for group 0 vs. group 1 and p-values for a significant difference in distribution, for each of the top features found by the Stabl algorithm. All features are from the CyTOF phenotyping assay.

### 3.4 Predictive Modeling using Stabl features

To test the top features found by Stabl, the three features that were found on the same random seed were taken (CD161+ CD45RA+ Tregs, CD14+ CD33+ monocytes, and NKT cells). An overall pipeline was created with the preprocessing used earlier, feature selection determined by the features found by Stabl, and a final model. The final model was a Logistic Regression with ElasticNet penalty, saga solver, balanced class weights, and max iterations of 1*e^6^. The model described in the pipeline was fitted to the training data, and the ROC curves were displayed for both the training and testing data, as shown in Figure 6. The Stabl features achieved an AUROC of 0.673 (95% CI of 0.553-0.774) on the training data, and 0.634 (95% CI of 0.432-0.823) on the testing data. Stabl employed bootstrapping to calculate the confidence interval, repeatedly resampling the data with replacement and using the distribution of the AUROC across 1,000 iterations to estimate it.

**Figure 6.**
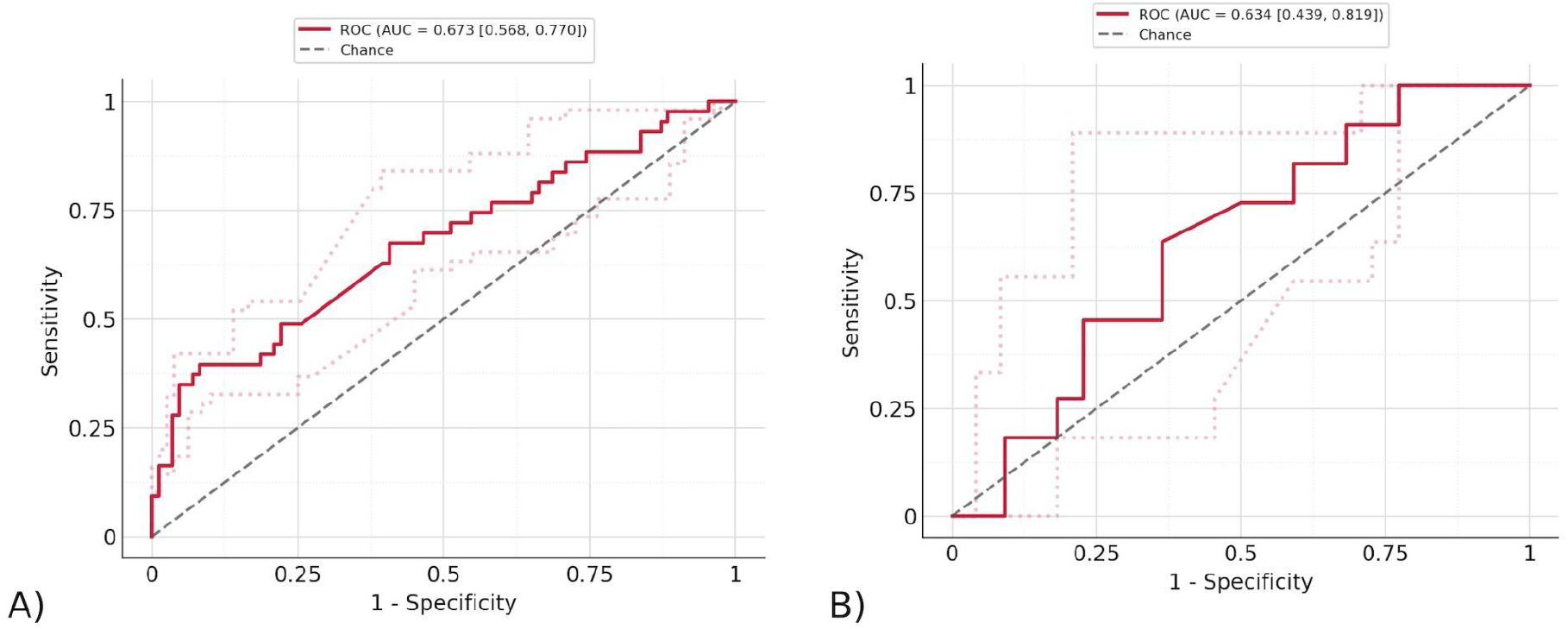
The ROC curve of the model trained by Stabl features - **A)** The ROC curve on testing data. The AUROC of the testing data using exclusively the features found by Stabl is 0.673 (95% CI of 0.553-0.774) and training data. **B)** The ROC curve on training data. The AUROC of the training data using exclusively the features found by Stabl is 0.634 (95% CI of 0.432-0.823).

## 4. Discussion and Conclusion

Overall, this project proved useful in exploring MOFA and Stabl as complementary algorithms for exploring the underlying trends in heterogeneous data collected across years and from different sources like the FluPRINT dataset. We demonstrate the utility of MOFA and Stabl in discovering the principal features that correlate with high vs. low influenza vaccine responders. By applying both unsupervised (MOFA) and supervised (Stabl) approaches to comprehensively analyze the FluPRINT dataset, we identify novel biomarkers through complementary analytical perspectives.

As an unsupervised model that is not specifically looking for top features for outcome, MOFA was used to examine the structure of the data and find if there were any connections or similarities between the top features for decomposition versus outcome. MOFA’s interpretability and various graphs made it easier to view the connections between factors, features, group, and the data as a whole. Due to its exclusion of missing values in likelihood estimations when performing the decomposition into the latent factors, it was especially useful for FluPRINT due to the high portions of missing values as there was no need for imputation and then chance of bias that could then throw off the model.

Meanwhile, Stabl was used to find the top features across the entire dataset that were most significant in predicting the flu vaccine response, rather than in estimating the population structure. Stabl’s thorough model produced results that allowed for better visibility of the top feature’s performance on a general scale and compared to other features, giving a better understanding of the significance (measured through p-value) and accuracy (measured through AUROC of predictive models) of the features found. It also had a higher AUROC (0.673 - 95% CI of 0.432 to 0.823, 0.634 – 95% CI of 0.553 to 0.774) as compared to MOFA (0.616, 0.616) on both training and testing data. This is due to being designed specifically for feature selection and using a supervised approach, while top features were instead extracted from MOFA’s decomposition of the data, which was found through unsupervised machine learning methods. The higher AUROC achieved with Stabl’s features are likely due to the advantage of knowing the outcome, which was discrete and easily defined through a binary outcome, in a dataset that otherwise has considerable noise because of missing values.

The Stabl algorithm selects features based on their stability and predictive value across multiple bootstrap resamples and regularization paths, rather than solely on individual statistical significance (p-values). Thus, it is possible for certain features to be consistently selected by Stabl due to their contribution to the predictive performance of the model when considered jointly with other features, even if individually they do not show strong statistical significance when tested independently. In other words, these features may provide complementary predictive information when combined with other selected features, improving the overall predictive accuracy of the model. Also, the relatively small sample size and inherent variability in the data may limit the statistical power to detect significant differences for some individual features.

In explaining the structure of the data, MOFA found many features with highly significant p-values (calculated through a t-test on the two distributions of the value of the features (i.e. the group 0 and group 1 samples)), including IL neg 2 CD4 pos CD45RA neg pSTAT5 and IL neg 2 CD8 pos CD45RA neg pSTAT5. The importance of these features has been suggested by recent research, which indicated that a subset of effector memory CD8 T cells produce IL-2, whose signaling contributes to protection against a chronic viral challenge (15). These results were not similar to the features of Stabl or the results of the previously published SIMON study. However, this is expected due to MOFA being designed to serve a purpose similar to principal component analysis rather than outcome prediction. Since the algorithm does not directly search for features that accurately predict response, these features had to be isolated using the correlation between factors and outcome, followed then by taking the most weighted features for the most highly correlated factor. This provides a likely explanation as to why MOFA’s results differ from those of Stabl and SIMON.

On the other hand, Stabl’s selected group of features proved to be effective in predicting response. All results were T cells (either CD4+ or CD8+ subsets), expected due to their key role in boosting immunity by recognizing and targeting conserved epitopes of similar viruses (16). Some of the results corroborated by the SIMON study, proving the validity of the algorithm, which also found novel features. Overall, the top features found by Stabl were: CD161+ CD45RA+ Tregs, CD14+ CD33+ monocytes, NKT cells, and CXCR5+ CD8+ TFH T cells, which were all higher in high responders.

The most significant feature found by Stabl was CD161+ CD45RA+ Tregs, whose validity was proved as it appeared in multiple trials with many different random seeds. It was supported by the SIMON algorithm which found the same feature as most important. This indicates the likelihood of this feature to be a top indicator of vaccine response, especially as they are IL-17A producing memory cells that have proved to be beneficial in other pathogenic conditions. It has also previously been associated with the influenza virus in mice (17).

Our t-test analysis for individual features confirmed Stabl’s top finding, with CD161+ CD45RA+ Tregs showing the most significant difference between high and low responders (p < 0.05). This concordance between simple univariate statistics and our more sophisticated algorithm validates the robustness of this particular feature. However, while t-tests can identify individual differentially expressed features, they’re fundamentally limited when applied to multi-omics datasets with thousands of features due to multiple testing issues and inability to capture feature interactions. Stabl provides critical advantages for multi-omics analysis through its stability selection framework, which protects against spurious correlations through bootstrapping and artificial feature injection. While simple statistical tests may identify key individual markers, algorithmic approaches like Stabl are necessary to build robust predictive models from high-dimensional data while controlling false discovery rates. The AUROC of the predictive model using just the CD161+ CD45RA+ Tregs was 0.563 (95% CI of 0.412-0.703), lower than that of the model with Stabl’s top features, which includes the other features it found as well.

Stabl also found CD14-CD33-CD3+ CD4-CD8+ Non-naive CD8+ CXCR5+ TFH T cells, which were found to share a parent population with a feature from the previously published SIMON study and has previously been connected to viral infections by influencing antibody response through B cell interactions. The other two cell subsets it highlighted included CD14+ CD33+ classical monocytes and NKT cells. Monocytes can be important for their ability to recognize antibody-bound target cells via Fc receptors (18). Similarly, NKT cells have direct correlations to better immune response through their release of multiple cytokines and chemokines that boost adaptive responses (19). Although the results prove promising, there is considerable uncertainty, as the models may be random since the AUROC confidence intervals fall below 0.5.

Still, MOFA and Stabl were extremely beneficial in this deeper examination of FluPRINT, displaying their potential to contribute to progress by providing an easier, accurate way of understanding data. In conclusion, these algorithms helped to gain valuable knowledge about the underlying population structure of this flu vaccine dataset by working together to explain the biological reasons behind the makeup of the existing data collected from large populations and taking a step forward in working towards accurate future prediction of response.

## Supplementary Table 1 Acknowledgements

We are very grateful to Tyson Holmes for detailed review of an early draft of the manuscript and for helpful initial statistical guidance.

